# Prediction of nanographene binding-scores to trout cellular receptors and cytochromes

**DOI:** 10.1101/2021.02.20.432107

**Authors:** M. C. Connolly, J.M. Navas, J. Coll

## Abstract

To address the increasing concerns surrounding possible impacts of graphene-related materials on the aquatic environment, this study focused on computational predictions of binding between models of graphenes in the nm size range (nanographenes, nGs) and the aryl-hydrocarbon receptor (tAHR) and P450 cytochromes (tCYPs) of rainbow trout (*Oncorhynchus mykiss*). The tAHR plays a key role in the induction of detoxifying and early immune responses and tCYPs are essential for detoxifying planar hydrophobic chemicals such as nGs. After 3D modelling of those trout proteins, docking algorithms predicted the size-dependance profiles of nGs binding-scores to tAHR and tCYPs in the low nM range (high binding-affinities). Virtual oxidations of nGs to nGOs (carboxy-, epoxy-and/or hydroxy-oxidations) further lowered the corresponding binding-scores in level/type-oxidation manners. Among all the tCYPs, the tCYP3AR (the equivalent to human CYP3A4) was identified as a potential key interaction enzyme for nGs because of its lower binding-scores. These results implicate a possible processing pathway to be further probed through *in vitro* and *in vivo* experimentation. Together the information generated can be pivotal for the design of safer graphene-related materials for a variety of applications and help to understand their detoxification in aquatic vertebrates.

## Introduction

Graphene-related materials (GRMs) have potential applications in a variety of human activities, from aeronautics to biomedicine, to electronics and in agriculture. This is causing a constant increase in the production of GRMs that will likely lead to releases to the environment during production, use and/or disposal ^1^. Although the current concentrations and toxicological hazards of GRM in the aquatic media are yet minimal, they are expected to increase with graphene production ^2-4^. Usually, graphene is not used in its pristine form but with a certain degree of oxidations (GOs). GOs have been shown to interact with other environmental pollutants, such as polyaromatic hydrocarbons (PAHs) causing an increase in cytotoxicity as estimated in fish cell lines ^5^. Once in the aquatic environment, residues of GOs will likely interact with a variety of biota but there is only a limited knowledge of the bio-reactions they might induce. Among these bio-reactions, perturbations in detoxifying mechanisms ^6^ and immune responses ^7-9^ would have important impacts on vertebrates. To better understand the possible initiation of detoxyfying bio-reactions induced by or interfering with GOs, we have selected small fragments or nanographenes (nGs) to mimic degradation products of larger sheets. In this study we have focused on the possible effects of pristine nanographenes considering those as a single layer sheet < 100 nm lateral size without (nGs) and with different levels of oxidation (nGOs).

Previous work has shown that nGOs can adhere to the cellular plasma membrane, insert into the lipid bilayers and enter the cytoplasm ^10^. Once inside the cell, detoxification pathways could be activated that may hypothetically include receptor binding and induction of enzymes that cause a reduction of hydrophobicity by further oxidation (including carboxy-epoxy-and hydroxy-oxidation) to more hydrophilic metabolites which could more easily be excreted via the bile, urine or gills to prevent toxic concentrations and their derived bio-effects ^11, 12^. We hypothesize that extracellularly, large graphenes (platelets) may begin their degradation by different mechanisms such as oxidation with reactive oxygen produced by peroxidases ^13, 14^, leading to nGO. Alternatively, and although not yet demonstrated, different intramolecular imperfections in the graphene platelets could allow oxidations to occur extracellularly. Once inside cells, nGOs could bind to the aryl-hydrocarbon receptor (AHR) and induce oxidating cytochromes (CYPs), that will feed-back to further accelerate oxidation.

Taking into account the hypothesis mentioned above, we have computationally explored whether nGs/nGOs of ∼25-200 rings could bind to intracellular AHR and/or CYPs with enough binding affinity to explain any downstream effects. Given the large differences among nGOs derivatives, it would be most convenient to select only those with the highest likelihood to elicit such bio-reactions that could be then demonstrated through *in vitro* or *in vivo* experimentation. Any reduction in the number of possibilities would make experimentation approaches more targeted and feasible. Therefore, to improve the likelihood of experimental success and to increase our understanding of possible detoxification mechanisms, *in silico* binding predictions between nGs/nGOs and 3D modeled AHR / CYPs has been performed.

For this work, rainbow trout (*Oncorhynchus mykiss*) was selected as a model vertebrate species because of their high sensitivity to environmental pollutants, and widespread use in ecotoxicological studies. Additionally, a number of their detoxifying and immune-related genes and/or functionalities have been studied. Therefore, this work specifically focuses on the trout aryl-hydrocarbon receptor (tAHR) and detoxification cytochrome enzymes such as CYPs (tCYPs).

Single layer graphene is constituted by a series of aromatic rings forming a planar structure. Therefore, Gs/GOs are structurally related to PAHs which are also constituted by aromatic rings albeit PAHs have a much lower number of rings. Since the AHR is one of the first proteins to recognise the intracellular presence of PAHs, we have hypothesized that the AHR could also bind nGs/nGOs. Once bound to a ligand, the cytoplasmic AHR dissociates from HSP90 and other proteins, binds to the aryl-hydrocarbon receptor nuclear translocator protein (ARNT) ^15^ and translocates into the cell’s nucleus. The co-transcriptional factor AHR/ARNT complex then interacts with xenobiotic genomic DNA response elements (XREs) to induce transcription of *cyps* and many other proteins ^16^. The recognition of ligands by PAHs has been mapped in mammalian AHRs to their PAS domain (<u>P</u>ER, <u>A</u>RNT, <u>S</u>IM) which is common to other proteins of the same family of transcriptional factors. In particular, deletion analysis demonstrated that the binding-domain was localized in the amino-terminal part of AHR at residues 1-403. In contrast, deletion of 398-805 residues had no effect on ligand binding ^17, 18^. Mouse AHR mutants at the 375 residue confirmed a reduction of binding of the prototypical ligands ^19^. Although all the above mentioned properties are most probably similar in trout, confirmatory studies are very scarce and some AHR downstream pathways may differ ^20^.

Detoxification of target aromatic molecules like PAHs, begins with the so called detoxification phase 1 by the addition of -COOH, -O, or -OH, by monooxygenases (predominant), flavoprotein monoxygenases, monoamine oxidases, epoxide hydrolases and/or reductases. Among all these possibilities, CYPs were chosen for this study because in vertebrates these oxidative microsomal-membrane heme-containing enzymes are the main participants in phase 1 detoxyfication ^21^. In addition, some of them are induced after ligand binding to AHR and have been studied in rainbow trout ^22^, to such an extent that induction of CYP1 by AHR activation is been used as a biomarker for hydrophobic contaminant exposure ^23, 24^. A feed-back loop regulates AHR and CYPs transcription, increasing CYP1 down-regulates *ahr* and inhibition of CYP1 up-regulates *ahr* ^16^. In human cells, inhibition of CYP1 up-regulates also helper T cells (and other immune cells) while down-regulating *il17* gene expression ^25^. Furthermore, some of the CYPs effects during infections resemble those of proinflammatory cytokines, further implicating CYPs in inflammation and immunology ^26^.

Putative CYPs predicted in their corresponding genomes (http://drnelson.uthsc.edu/cytochromeP450.html), showed species-and family-specific amino acid sequences ^22^, nevertheless with high tridimensional 3D structural similarities. CYPs are divided into those involved in xenobiotic catabolism (CYP1-4) and those implicated in metabolic biosynthesis (CYP5-51). Some CYPs are constitutively expressed but others are induced by PAHs ^27^. Specifically in fish, many CYP enzymes are known to be implicated in fish metabolic biosynthesis such as that of hydroxycholesterols ^28^, but also several CYP1-4 detoxyfying enzyme families have been detected ^22^. Therefore, it is expected that most foreign compounds introduced into the cellular environment of a fish body, including nGs/nGOs, could hypothetically interact with several CYPs as well as with other detoxifying enzymes. Furthermore, structurally distinct nGs/nGOs may have specific binding affinities with different CYPs with either inhibitory or synergistic results.

Therefore, this work focuses on prediction of binding-scores and conformations or binding-poses of pristine nGs and oxidated nGOs with tAHR and tCYPs. To our knowledge, no such computational predictions have been yet reported for any graphene-like or nGs/nGOs to tAHR/tCYPs ^6, 29-31^.

## Materials and Methods

### Nanographene ligands

Graphene-like ligands were obtained by a similarity search in PubMed (https://pubchem.ncbi.nlm.nih.gov/#query=C1%5BC%40%40H%5D2N%3DN%5BC%40%40H%5D3%5BC%40H%5D1C1%3DC(C%3DCC%3DC1)%5BC%40H%5D23&tab=similarity) by providing an 8 ring graphene-like molecule (SMILES= C1=CC2=C3C=CC4=CC=C5C=CC=C6C7=CC=C8C=CC(=C1)C2=C8C7=C3C4= C56) corresponding to Zinc ID 104923703 (http://zinc15.docking.org/). The corresponding downloaded sdf file manually curated, contained 86 small molecular weight graphene-like compounds.

To mimic graphene platelet degradation products, pdb models of nanographene (called nGs here, because of their 10-200 nm lateral side range) were designed and obtained using the open-source Phyton GOPY tool (https://github.com/Iourarum/GOPY). The nGs were obtained in different sizes (4 to 156 rings), and with different levels of oxidation (carboxylation -COOH, epoxy oxidation -O-, hydroxylation –OH and/or combinations). Oxygens were added to a maximum of ∼1.6 Carbon/Oxygen C/O ratio, similar to those described in previous experimental work ^32^. While carboxylation can occur only at the C edges, both epoxyoxidation and hydroxilation may occur at each of the Cs inside the rings to be distributed randomly between the two faces of the planar nG sheet (see some depictions in Figure S3). In all cases, steric hindrance may reduce the number of theoretical possibilities. Ten random designs for each of the oxidation scenarios were computationally constructed for each oxidation state by modifications of the Phyton GOPY tool, which took into account all the requirements mentioned above^33^. The resulting nanographene structures visualized in PyMOL showed a variety of numbers and intramolecular locations approximating the input numbers provided to the GOPY tool because of steric restrictions. Additionally, docking analysis rejected 10-20% of the structures specifically when 2 oxidations were too close in the same ring.

### Tridimensional trout AHR and CYP 3D models

The trout AHR (tAHR) was modeled using the SWISS-MODEL homology modeling (https://swissmodel.expasy.org/interactive). Structural similarities were expressed in Angstroms Å / number of common atoms, as estimated by 3D superposition with hAHR-1 in the CCP4 Molecular Graphics program vs2.10.11 (http://www.ccp4.ac.uk/MG) (Table 1)

**Table 1.** Modelling of human and trout AHR

| name | protein sequence | #aa | SWISS template | #aa 3D modeled | SI, % | RMSD, Å/#atoms |
| --- | --- | --- | --- | --- | --- | --- |
| hAHR | NM_001621 | 848 | 4m4x | 107-261 | 100 | 1.92/103 |
| hAHR-1 | NM_001621 | 848 | 5sys5 | 34-391 | 28.7 | 0.00/360 |
| hAHR-2 | NM_001621 | 848 | 4f3l | 35-391 | 24.7 | 1.27/99 |
| hAHR-3 | NM_001621 | 848 | 4f3l | 29-381 | 24.8 | 2.74/122 |
| hAHR-4 | NM_001621 | 848 | 5sy5 | 36-381 | 21.2 | 1.88/115 |
| hAHR-5 | NM_001621 | 848 | 4zpk | 32-386 | 29.5 | 1.99/258 |
| hAHR-6 | NM_001621 | 848 | 4f3l | 36-381 | 26.6 | 1.17/95 |
| hAHR-7 | NM_001621 | 848 | 5sy7 | 33-373 | 26.4 | 5.48/104 |
| tAHRa | AF065137 | 1058 | hAHR-1 | 34-391 | 71.2 | 0.38/348 |
| tAHRb | AF065138 | 1059 | hAHR-1 | 34-391 | 71.2 | 0.38/348 |

The sequences were obtained from the NUCORE databank. Protein sequences from PAS domain of the hAHR (residues 1 to 420) were submitted to the SWISS-MODEL homology modelling, which automatically selected 7 possible templates with the highest sequence identity (SI). The hAHR-1 (highlighted grey background) was selected as model template for tAHRa and b. tAHRa and b sequences showed 97 % SI. hAHR, human AHR.

The trout CYP (tCYP) enzymes selected for this work corresponded to those sequence-reviewed, that were available for *Oncorynchus mykiss* and downloaded from UNIPROT (https://www.uniprot.org/uniprot/?query=reviewed:yes). They corresponded to most of the best studied tCYPs ^22^. Human eosinophil peroxidase or myeloperoxidase, was also included for comparison with previous work because to our knowledge it was the only CYP enzyme used in a previous nG oxidation computational study ^34^. The amino acid sequences were submitted to the SWISS-MODEL to obtain hypothetical 3D models. The coordinates of the heme and ligand binding pockets provided by the CYP human templates were copied from the human pdb to the modeled trout pdb file (**R**esearch **C**ollaboratory for **S**tructural **B**ioinformatics, RCSB, **P**rotein **D**ata **B**ank PDB). All modeled structures were visualized in PyMOL and PyRx. Structural similarities were studied as commented above by 3D superposition with tCYP1A1.Predicted binding pockets and α-helices were investigated using the seeSAR vs.10 program (https://www.biosolveit.de/SeeSAR/).

### AutoDock Vina virtual docking

The AutoDock Vina program ^35^ included in the PyRx 0.9.8. package ^36^ (https://pyrx.sourceforge.io/) was used in e7 desk computers to predict Gibbs free-energy (ΔG) as described before ^28, 37, 38^, using grids including the whole protein molecules (blind docking) or the identified binding pockets when defined in the CYP templates. The *.sdf files were converted to *.pdbqt files by allowing ligand rotatable bonds, adding protonation and adding Gasteiger−Marsili partial atomic charges with one ffu energy minimization step carried out by the Open Babel program included into the PyRx package. Water molecules were not considered. Only the binding-poses with the lowest binding energy of each *.out.pdbqt were retained for further analysis. The output ΔG energies were converted to constant inhibition (Ki) values in molar concentrations (M), using the formula Ki = exp([ΔG × 1000] / [R × T]) (R = 1.98 cal/mol, and T = 298 °C)^39^. Final values were converted to nM. The predicted structures were visualized in PyRx and/or PyMOL (https://www.pymol.org/).

### SeeSAR modeling

The seeSAR seeSAR vs.10 package (https://www.biosolveit.de/SeeSAR/) ^40-42^ was used for binding pocket predictions and alternative visualization of bound complexes as described previously^28, 37, 38^

## Results

### 3D Modeling of trout AHR

One of the requirements for this study was to obtain a 3D model of the PAS domain of trout AHR (tAHR) based on the two amino acid sequences of tAHR available (97% identical) and some corresponding 3D solved structures. Since only residues 107-261 of human AHR (hAHR) were crystallographically solved and those were not enough to cover all the corresponding hAHR ligand binding domains, a template search was undertaken for a better template. Seven hAHR models were selected by the SWISS-model program using template PAS domains from diverse proteins including those from similar transcription factors. Among them, 6 models showed < 2 Å RMSD differences in their 3D structure alpha-carbons. The hAHR-1 model covering the highest number of amino acids of 357 residues was selected (Table 1, highlighted in grey background) to model the two tAHRs. The resulting modeled tAHRa and b structures were very similar (Table 1, last entries).

### 3D Modeling of trout CYPs

Because of the absence of crystallization studies that solved the structures of trout CYPs (tCYPs), their hypothetical 3D structures were modeled for the first time (Table 2, Figure S1). Despite the differences in their amino acid sequences (Table S1), all the tCYP models have similar overall 3D structures as shown by their < 3 Å RMSD. The tCYP 3D structures have several α-helices, one conserved Cysteine, and an inner heme. Their carboxy-terminal helices contain a highly conserved domain (Table S1, red amino acids). Heme binding implicating residues ∼100-140 and ∼300-320 were in the pM range, as shown by seeSAR. An amino-terminal tunnel-like access (Figure S1, grey arrow) may be the main route for substrates/inhibitors to penetrate nearby the heme for optimal oxidation. A binding pocket predicted by seeSAR, was identified around the heme in all tCYPs (Figure S1, yellow pocket).

**Table 2.**
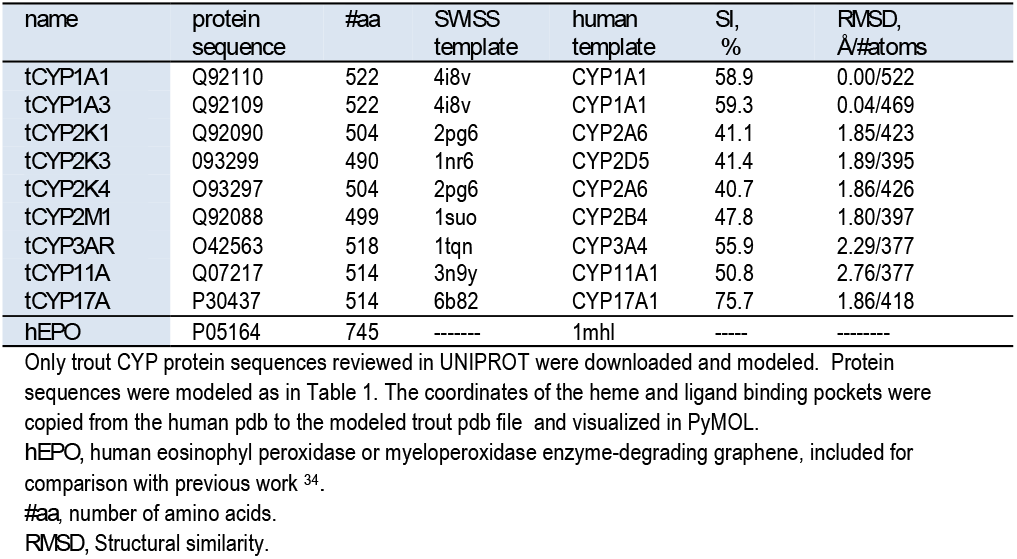
Modelling of trout CYPs compared to human myeloperoxidase hEPO

### Computational binding of nG-like compounds to hAHR and tAHR

Most but not all nG-like compounds showed similar binding-scores to h and tAHR despite their differences in amino acid sequences (Figure 1A). Nevertheless, their corresponding nG binding-poses were different (Figure 1B and C).

**Figure 1.**
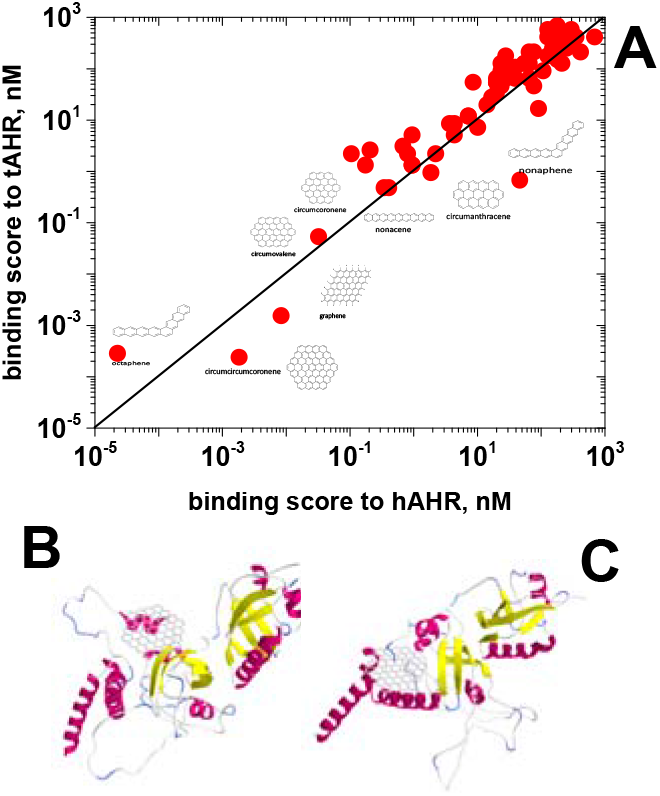
Binding-scores of nG-like compounds to hAHR and tAHR. **A)** Binding-scores to 83 nG-like compounds compared between hAHR and tAHR. **B)** Drawing showing the best pose of 37-ring nG bound to hAHR. **C)** Drawing showing the best pose of 37-ring nG bound to tAHR.

Among the nG-like compounds there was a reduction of binding-scores down to 10^−4^ nM with increasing number of rings per molecule, in the following order, circumanthracene (13 rings), circumcoronene (19), circumvalene (24), Ng (25) and circumcircumcoronene (37).

Since there were no differences between the binding-scores of tAHRa and b (not shown), the tAHRa model was selected for the rest of the work. Our approach predicted binding-scores in the 10^3^ nM range for the hAHR prototypical agonist 2,3,7,8-tetrachlorodibenzo-para-dioxin (TCDD) (not shown).

Modeling of 20×20 nm nGs resulted in nGs of 72 rings down to binding-scores of >10^−10^ nM, compared to those of 25 nG or 37 circumcircumcoronene rings which bound to the hAHR / tAHR with binding-scores in the ∼ 10^−3^ nM range (Figure 1A). To further investigate the influence of nG size in the binding to tAHR and tCYPs, different sizes were designed and computationally tested by blind docking (PAS tAHR domain) or binding-pocket docking (tCYPs).

The results predicted that the lower binding-scores to tAHR corresponded to nGs > 100-rings. Larger nGs maintained these binding-scores. Binding to tCYPs showed a similar size-profiles as to tAHR, but in the 10^−5^-10^−9^ binding-score ranges (Figure 2, blue, green and red lines). Most tAHR binding to nGs mapped to its PAS ligand binding site. In contrast, most nGs binding to CYPs were localized on their surfaces away from their heme sites, suggesting some unspecific bindings (Figure S2). Only 4-9-ring nGs were mapped nearby to their heme site, most probably because only such small compounds could penetrate through the CYP tunnels to such internal location to be specifically oxidized (Figure S2).

**Figure 2.**
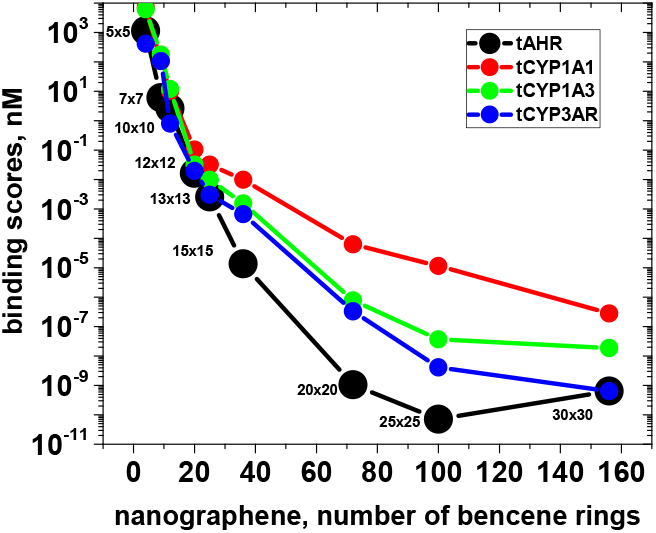
Influence of the number of rings of nG in their binding to tAHR and to tCYPs. nGs of different sizes were designed by the GOPY tool. **Black numbers**, lateral sizes in nm of the nGs for inputs to the GOPY tool. **Black circles**, tAHR. **Red circles**, tCYP1A1. **Green circles**, tCYP1A3. **Blue circles**, tCYP3AR.

Models of 9 (7×7 nm, 32 Carbons) and 25 (13×13 nm, 77 Carbons)-ring nGs were selected for further studies, because significant binding-scores could be obtained in minimal computational times.

### Computational binding of nGOs to tAHR

Carboxy (**-COOH**), epoxy (**-O-**) and hydroxy (**–HO**) oxidations were mimicked by adding them to the 9 and 25-ring nGs to randomly generate their corresponding nGOs using 1.6 Carbon/Oxygen C/O ratios ^10^ (see some of their structures in Figure S3).

In this test, docking conditions were adjusted to yield binding-scores of ∼10^−3^ nM and ∼10^1^ nM for 25- and 9-rings nGs, respectively. Results with nGOs predicted 10^1^-10^−7^ nM binding-scores for 25-rings (Figure 3,) and 10^0^ nM for most 9-rings (Figure 3, red- and grey-edged circles, respectively). In particular, epoxidation (-O-) alone or in combination with carboxylation (-COOH), was the most efficient process to reduce the binding-scores of 25-ring nGOs. In contrast, hydroxylation (-OH), and its combinations increased their binding-scores.

**Figure 3.**
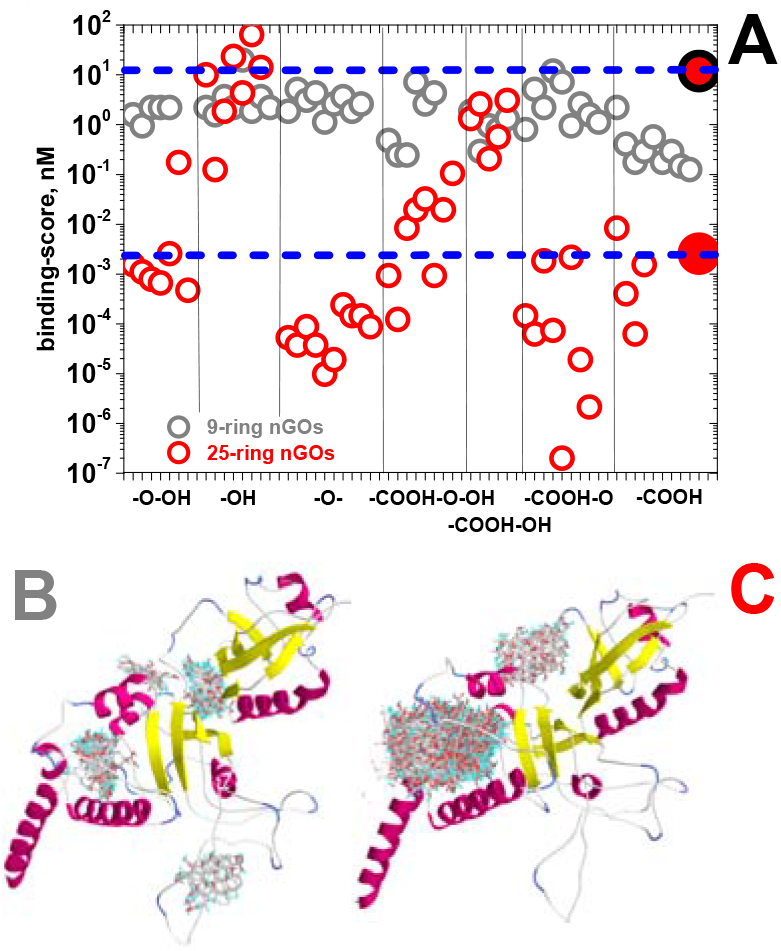
Influence of nG oxidation(nGO) in the binding-scores to tAHR. Oxidations with –COOH, -O-, -OH and combinations of them were randomly added at maximal C/O ∼1.6 ratios to 9 or 25-ring nG using the GOPY tool. **A) Grey-edged circles**, 9-ring nGO. **Red-edged circles**, 25-ring nGO. **Grey-edge, red filled circle**, 9-ring nG. **Red-edge red filled circle**, 25-ring nG. **Black vertical lines**, separation between different types of oxidations. **Dashed horizontal blue lines**, nGs binding-scores to tAHR. **B)** Mapping of 9-ring nGOs bindings to tAHR. **C)** Mapping of 25-ring nGOs bindings o tAHR.

Mapping of the corresponding 3D binding-poses predicted that the amino terminal end of the PAS domain of tAHR was targeted by most 25- and 9-rings nGOs (Figure 3C and B, respectively).

### Computational binding of nGOs with different oxidation levels

To study the influence of the levels of nGO oxidation on the binding to tAHR, different numbers of oxygen-containing molecules were added to 25-ring nGs. Results showed that increasing the level of peroxidation (**-O-**) to 70 % caused a continuous reduction to 10^−4^ nM binding-scores (Figure 4, green circles). Increasing the level of carboxylation (**-COOH**) or hydroxilation (**-OH**) to <25 % reduced their binding-scores to ∼10^−5^-10^−6^ nM, but increased them at higher oxidation levels (Figure 4, red and blue circles). Both nG and most nGO binding-scores to tAHR remained < 10^1^ nM, which may be significant compared to the 10^3^ nM binding-score of the AHR prototypical agonist TCDD under the same docking conditions.

**Figure 4.**
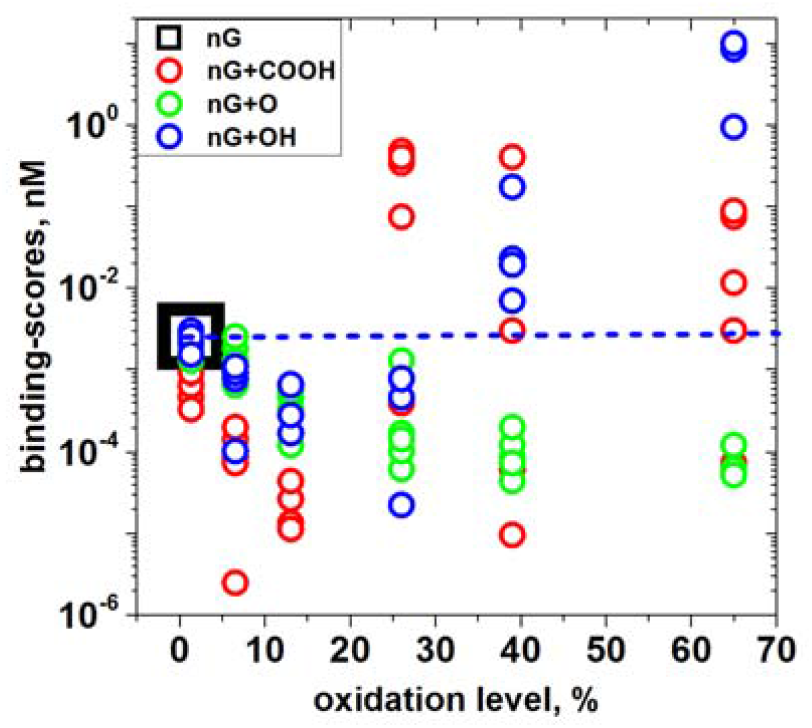
Influence of the level of oxidation in the binding of nGs to tAHR. Individual **–COOH, -O-, -OH** oxidant molecules were randomly added into 25 rings (77 Carbons) nGs by the GOPY tool. From 5-10 nG molecules were virtually oxidated per each level of oxidation. Due to steric problems when > 20-30 atoms were added, some of the resulting nGO structures when examined in PyMOL, only approximately confirmed the initial numbers supplied to the tool. Theoretical oxidations per nG molecule were expressed in % as calculated by the formula, 100 x number of output oxidative molecules per 77 Carbons. **Black big square and blue-dashed horizontal line**, binding-scores of 5×5 nm nGs (mean of n=5). red **circles**, nG+COOH. **Green circles**, nG+O. **Blue circles**, nG+OH.

Because of the nGs tendency to bind non-specifically to tCYPs (Figure S2), the size of the nGs were reduced to 5×5 nm (4 rings) and grids of 25 x 30 x25 Å surrounding the tCYPs heme were used for docking.

Under the above mentioned conditions, the nG binding-scores varied for each tCYPs molecular species from 10^2^ to 10^4^ nM (Figure 5, **black**-dashed horizontal lines). By increasing the level of nGOs oxidation to ∼ 25 % most binding-scores decreased to different minimal levels for different tCYPs. However, increasing the level of most oxidations > 25 %, resulted in increased binding-scores for most tCYPs, except tCYP3AR. Thus, with >25 % oxidations, tCYP3AR binding-scores were further reduced to 10^−2^ by carboxylation (Figure 5, 3AR, red circles), and to a lower extent by peroxidation and hydroxylation.

**Figure 5.**
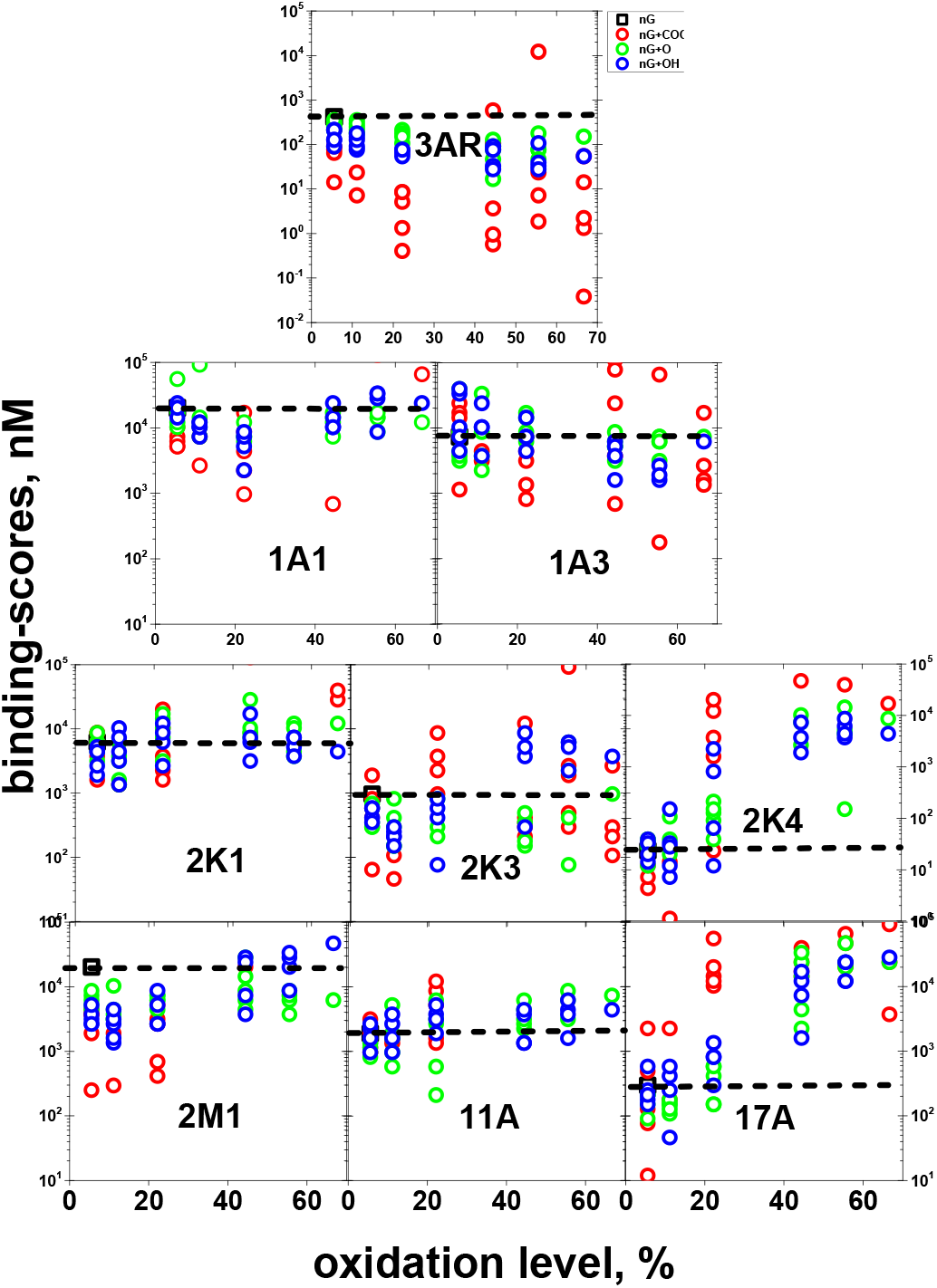
Influence of the level of oxidation in the binding of nGOs to tCYPs. Individual **–COOH, -O-, -OH** oxidant molecules were virtually and randomly added into 4 rings, 18 Cs nGs by the GOPY tool. From 5-10 nG molecules were virtually oxidated per each level of oxidation. Due to steric problems when > 20-30 atoms were added, some of the resulting structures examined in PyMOL only approximately confirmed the initial numbers supplied to the tool. Theoretical oxidations per nG molecule were expressed in % as calculated by the formula, 100 x number of oxidation molecules provided to the GOPY tool / 18 (n=5). The CYP family is in **black** bold lettering placed at around the 10^2^ nM label for easy reference. Only those nGs showing binding affinities <10^5^ nM were shown. All the scales were equal except the wider lower scale down to 10^−2^ for tCYP3AR. **Black big square and dashed horizontal line**, binding-scores of 5×5 nm nGs (mean of n=5). red **circles**, nGO-COOH. **Green circles**, nGO-O-. **Blue circles**, nG-OH.

## Discussion

Lateral sizes of commercial powered graphene are of ∼10 nm while the so called nanoplatelets range between 20-100 nm for oxidized graphene (GO) and between 100-1000 nm for carboxylated graphene ^10^. Our initial working hypothesis was that larger graphene fragments (e.g., platelets) are broken down and taken up by trout cells transformed into smaller graphene pieces of ∼10-100 rings (graphenes in the nm size range, here called nGs). We hypothesized that once internalized by cells, nGs/nGOs may interact with microsomal tAHR activating phase I detoxification mechanisms, such as those implicating peroxidase-dependent reactive oxygen ^13, 14^ and/or tCYPs enzymatic activities/*tcyp* gene transcription, to induce other bioeffects. All these possibilities remain largely unexplored.

To test such hypothesis by *in vitro* and *in vivo* experimentation, given the large numbers of different nGs/nGO derivatives, their numbers need to be reduced and possible candidates identified for further experimental studies. To do that, we proposed an *in silico* screening to select for those nGs with the highest binding affinities (lower binding-scores) to tAHR and tCYPs. We discovered that the maximal number of nG rings that tAHRs/tCYPs could bind is up to 25-rings with apparent binding-scores between 10^3^-10^−11^ nM and mapping to the PAS tAHR specific binding site defined by its prototypical TCDD ligand.

Increasing hydrophilicity that facilitates detoxification of graphenes, is mostly caused by carboxylation, peroxidation and/or hydroxylation, but ring-opening and reduction may also be implicated ^43^.Oxidation of nGs to nGOs by virtually adding -COOH, -O- and –OH such as those expected to be found in most graphenes used for biomedical applications, computationally predicted between 10-100-fold reduction in their nG binding-scores to tAHR depending on the numbers of oxidated carbons per molecule and the type of oxidation. Additionally, any possible implication of tCYPs in nG binding and subsequent oxidation may be sterically possible only with the smallest nGs according to the computational data. Predicted conformation or pose analysis showed that although nGs could bind tCYPs, most of those bindings seem to be unspecific or not specifically oxidative because of their excessive separation from the inner tCYP heme where oxidation is expected to be most active. Pose bindings nearby the tCYP heme could not be visualized for nGs > 4 rings. Only the minimal nGs of 4 rings were small enough to penetrate the inner heme located within the predicted binding pockets.

With such size restrictions, among all the CYPs, the tCYP3AR was predicted to bind stronger than other tCYPs to nGOs, specifically to those which were carboxylated at their edges. Perhaps the higher implication of tCYP3AR in additional oxidations compared to other tCYPs may be due to its participation in the metabolism of endogenous substrates, such as steroids ^43^. According to the UNIPROT data base, tCYP3AR is coded by the trout gene *cyp3a27*, however 3D modeling choose the human hCRP3A4 as the closest isomer, suggesting that despite nucleotide sequence differences, tCYP3AR is equivalent to hCYP3A4. Although our knowledge of any tCYPs is still scarce, immunologically relatedness of fish CYP3A to hCYT3A4 and rat CYP3A has been demonstrated ^44^. However, since there is evidence that tCYP3A4 shows no significant bio-effects to seven known substrates of the corresponding hCYP3A4 ^45^, the impact of substrate oxidation may differ among species. The hCYP3A subfamily is the most important of drug-metabolizing enzymes since it accounts for ∼30 % of the total CYPs in the liver and metabolizes ∼50 % of marketed drugs ^43^. In humans it is being expressed mainly in the liver where it accounts for 13 % of the total CYP content and has been implicated in the metabolism of ∼4% of marketed drugs ^43^. In contrast, few data are available for tCYP3AR.

The computational predictions identified the nGs molecular characteristics required to be suitable ligands for tAHR and therefore those with the higher potential to generate bio-effects. It may be hypothesized that subsequent translocation into the nucleus to induce the numerous tAHR-dependent genes would sustain the bases for a link between the environment and toxicological and immune responses at the cellular level. Although there are still no experimental evidences for that possibility, the present results indicate this may deserve futher explorations.

On the other hand, since vertebrate AHRs are expressed in many cells of both innate and adaptive immunity, the AHR may be one of the translators of environmental hydrophobic contamination into immunological responses. Experimental validation of such hypothesis, could be performed by *in vitro* and/or *vivo* assays by measuring the effect of synthetic nGs and their nGOs oxidated derivatives with different levels of oxidation, similar to those which have been investigated here using *in silico* screening.

Among the main difficulties to experimentally validate any hypothesis related to nG/nGO is the lack of such molecularly defined graphenes. An alternative to obtain the corresponding nGs/nGOs for experimental purposes could include not only chemical synthesis (if chemical synthesis could make it possible) but also strong sonication techniques or long peroxidation of graphene platelets. Techniques for removal of the largest fragments by ultracentrifugation and/or gel filtration chromatography are available. However, more detailed molecular characterization of such preparations than those routinely employed may also be required.

For further downstream experimentation, several *in vitro* assays such as those using subcellular fractions, fish cell lines, hepatocytes and/or tissues ^24^, could be used. Specifically to detect a variety of tAHR induced genes, for instance, RTqPCR could be used to detect up-/down-regulation of selected *tcyps*. Any induced tCYP1A and/or tCYP3A enzyme activities can also be monitored using ethoxyresorufin-O-deethylas(EROD) or benzyloxy-4-trifluoromethylcoumarin-O-debenzyloxylase (BFCOD) activity assays, respectively. To estimate possible induced immunological responses and/or to explain differences among AHR modulations observed across species^46^, concentration-dependent effects on trout immunosuppression ^25^, inflammation (downregulation of *il6, crp*, or upregulation of *tgfb*) and/or T cell and macrophage activation ^47^, may also be performed. Combining the information generated by *in silico* screening with future *in vitro* and *in vivo* experimentation, forms an integrative approach for unrevealling the bio-interactions and effects induced by nGs/nGOs and expanding our knowledge of these highly bio-interactive planar hydrophobic substances.

## SUPPORTING INFORMATION

**Table S1.**
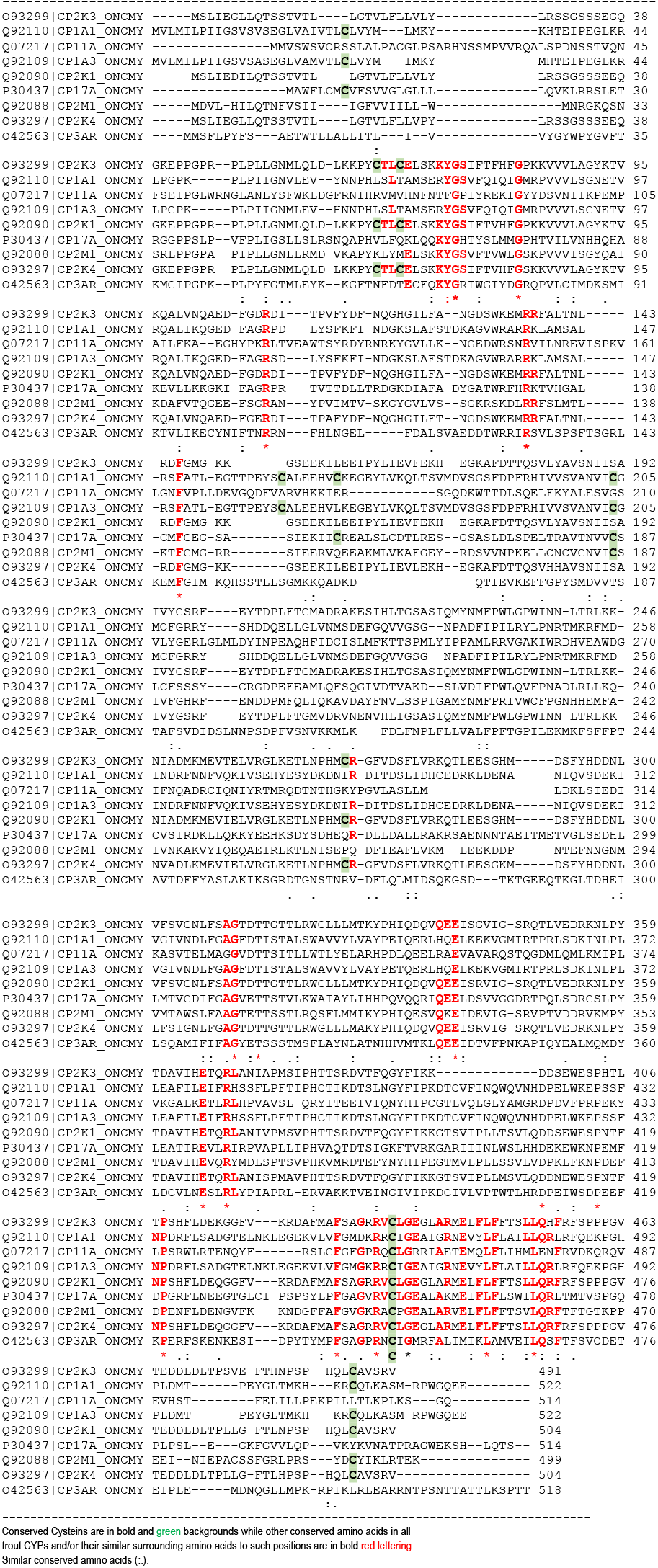
Trout CYPs CLUSTAL O(1.2.4) sequence alignment

**Figure S1.**
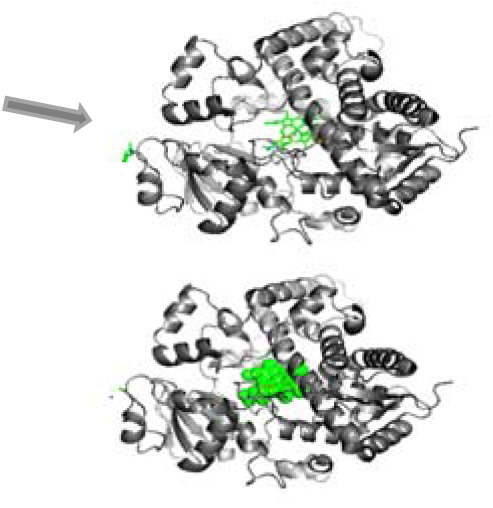
2D scheme of trout modelled CYP molecule and its predicted binding pockets. The figure shows the amino acid sequence in grey and the heme in green (up in lines and down in spheres) occupying an internal part of CYPs. A tunnel-like with an opening to the left in the drawing (Grey arrow) may support a possible way through which substrates and/or inhibitors may get close to the heme for optimal oxidation (ligand binding pocket). Most of the crystalized ligands map to that heme binding-pocket (yellow background). Between 6-15 other binding pockets were predicted by seeSAR depending on the CYPs (different color backgrounds in the down figure)

**Figure S2.**
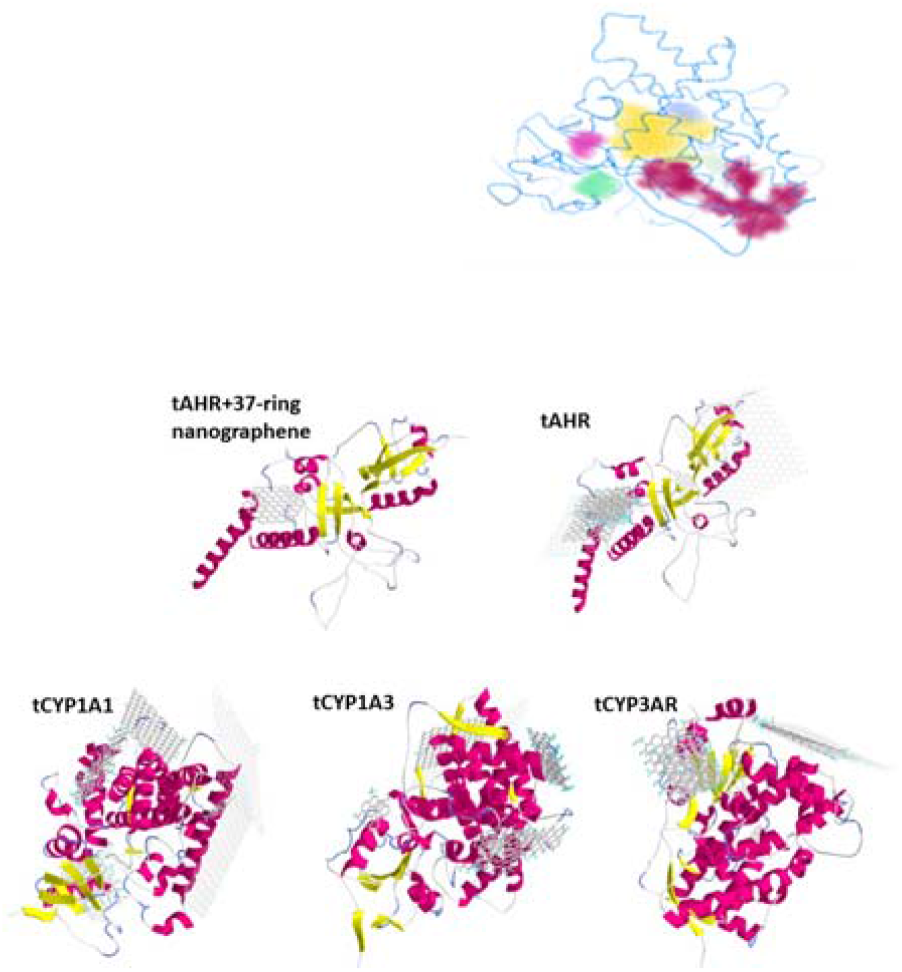
Drawings of nGs of different sizes bound to tAHR (up) and tCYPS (down). AutoDock Vina used a whole molecule grid (blind-docking). The nGs of 4 (5×5 nm), 9 (7×7 nm), 12 (10×10 nm), 20 (12×12 nm), 25 (13×13 nm), 36 (15×15 nm), 72 (20×20 nm), 100 (25×25 nm), and 156 (30×30 nm) rings were designed using the GOPY tool. Representative cartoons were made by combining one molecule of protein with several sizes of nG molecules and drawn in PyRx. The heme at the inner part of the CYP molecules have been removed for clarity.

**Figure S3.**
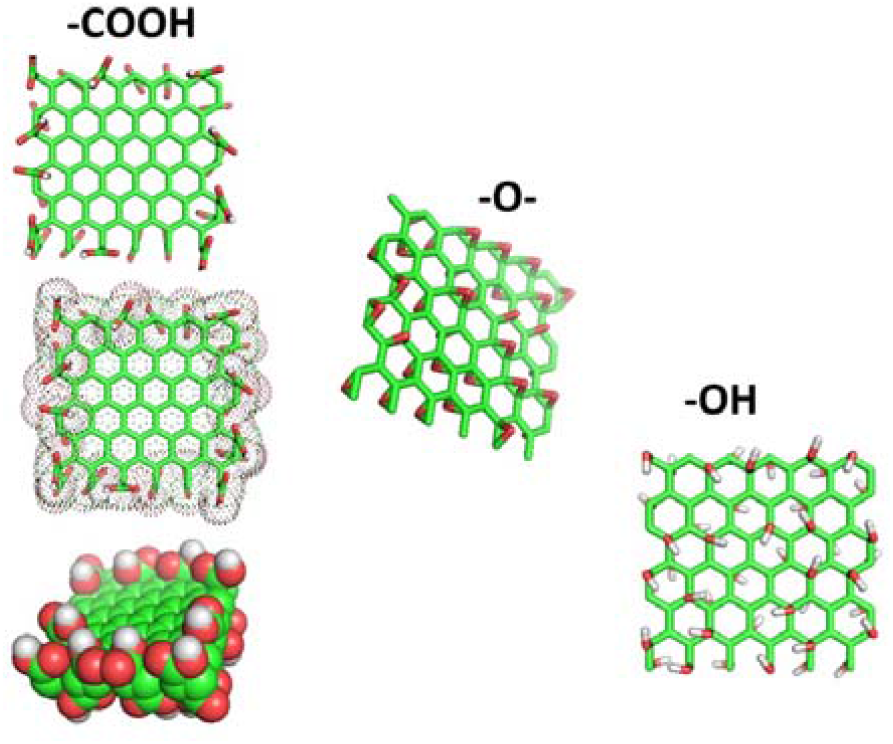
Cartoons of the nGOs of 25 rings (13×13 nm) designed by the GOPY tool. All the drawings were made in PyMOL.

## Authors’ contributions

MC and JN, collaborated by critically discussing strategies and writing and correcting the manuscript. JC performed and analyzed the computational work and drafted the manuscript. Authors read and approved the manuscript.

## Acknowledgements

Thanks are due to Dra Ana Valdehita for her help to obtain additional seeSAR licenses. Mona Connolly received financing granted by the Community of Madrid (2018-T2/AMB-11392, Mode 2, Young Doctor Recruiment).

